# Impacts on the structure-function relationship of SARS-CoV-2 spike by B.1.1.7 mutations

**DOI:** 10.1101/2021.05.11.443686

**Authors:** Tzu-Jing Yang, Pei-Yu Yu, Yuan-Chih Chang, Kang-Hao Liang, Hsian-Cheng Tso, Meng-Ru Ho, Wan-Yu Chen, Hsiu-Ting Lin, Han-Chung Wu, Shang-Te Danny Hsu

## Abstract

The UK variant of the severe acute respiratory syndrome coronavirus (SARS-CoV-2), known as B.1.1.7, harbors several point mutations and deletions on the spike (s) protein, which potentially alter its structural epitopes to evade host immunity while enhancing host receptor binding. Here we report the cryo-EM structures of the S protein of B.1.1.7 in its apo form and in the receptor ACE2-bound form. One or two of the three receptor binding domains (RBDs) were in the open conformation but no fully closed form was observed. In the ACE-bound form, all three RBDs were engaged in receptor binding. The B.1.1.7-specific A570D mutation introduced a salt bridge switch that could modulate the opening and closing of the RBD. Furthermore, the N501Y mutation in the RBD introduced a favorable π-π interaction manifested in enhanced ACE2 binding affinity. The N501Y mutation abolished the neutralization activity of one of the three potent neutralizing antibodies (nAbs). Cryo-EM showed that the cocktail of other two nAbs simultaneously bound to all three RBDs. Furthermore, the nAb cocktail synergistically neutralized different SARS-CoV-2 pseudovirus strains, including the B.1.1.7.

## Introduction

The emergence of the United Kingdom (UK) variant of the severe acute respiratory syndrome coronavirus (SARS-CoV-2), B.1.1.7, also known as the Variant of Concern (VOC) 202012/01 strain, was first identified in southeastern UK in December 2020^1^. B.1.1.7 was significantly more transmissible than the other SARS-CoV-2 variants across UK, with an estimated increase of the reproduction number, also known as R0, by 0.4 to 0.7^2^. B.1.1.7 quickly spread across the world, resulting in a surge of infected cases, and consequently stringent travel restrictions and national lockdowns in many countries. B.1.1.7 is defined by 23 genetic mutations compared to the original strain in Wuhan. In addition to the asparagine (D) to glycine (G) mutation at position 614 (D614G) of the surface spike (S) protein, which has dominated the reported COVID-19 cases since the summer of 2020 (GISIAD database: https://www.gisaid.org), B.1.1.7 harbors eight additional mutations within the S protein. Of particular interest is the asparagine (N) to tyrosine (Y) mutation at position 501 (N501Y), located at the receptor binding motif (RBM) that is involved in direct physical interactions with the human angiotensin conversion enzyme 2 (ACE2) to initiate viral entry into host cells^3,4^. While the functional implications of the remaining mutations remain to be established, the deletion of histidine 69 and valine 70 (ΔH69/V70) within the amino terminal domain (NTD) of the S protein is also present in the Danish SARS-CoV-2 mink-associated variant that caused zoonotic transmissions to infect humans, leading the culling of over 17 million minks in November 2020^5^. Although most neutralizing antibodies (nAbs) that have been identified from convalescent plasma mostly target the receptor binding domain (RBD) of the S protein to sterically prevent its engagement with ACE2, several nAbs were reported to target the NTD without blocking ACE2 binding^6–12^. In particular, tyrosine 144 of the S protein, which is deleted in B.1.1.7, is part of the structural epitope of the NTD-specific antibody 4A8. Indeed, a recent study shows that the ΔY144 deletion strongly diminishes the binding by multiple NTD-derived nAbs, including 4A8, by more than 1000-fold, suggesting that the increased infectiousness of B.1.1.7 is associated with its ability to evade host immunity^13^.

## Results

### Overall structure of SARS-CoV-2 S-UK (B.1.1.7)

We determined the cryo-electron microscopy (cryo-EM) structure of the ectodomain of the S protein of B.1.1.7 (hereafter S-UK) and identified four distinct conformations by three-dimensional variability (3DAV) analysis (Figure 1). Three of the conformation classes corresponded to one upward RBD (RBD-up) conformation (their nominal resolutions ranged between 3.2 and 3.6 Å), and the remaining one corresponded to two RBD-up conformation with a nominal resolution of 3.3 Å (Figure 1B and Supporting Information Fig. S1). Collectively, the one RBD-up conformations accounted for 73 % of the overall population and the two RBD-up conformation accounted for the remaining 27 %. Due to the intrinsic dynamics, the loop regions in the NTD where the ΔH69/V70 and ΔY144 lie, as well as the P681H mutation in the furin cleavage site could not be resolved with clarity in the cryo-EM maps. The remaining mutation sites, namely N501Y, A570D, D614G, T716I, S982A and D1118H, could be unambiguously resolved to build the atomic models accordingly (Supporting Information Fig. S2). Despite the large number of mutations, the overall structure of S-UK did not differ significantly from that of the original Wuhan strain and the D614G variant (hereafter S-WT and S-D614G, respectively), except for the relative populations and the number of the RBD-up conformation, which have direct implication in host recognition through binding to angiotensin converting enzyme 2 (ACE2)^4,14–20^.

**Figure 1.**
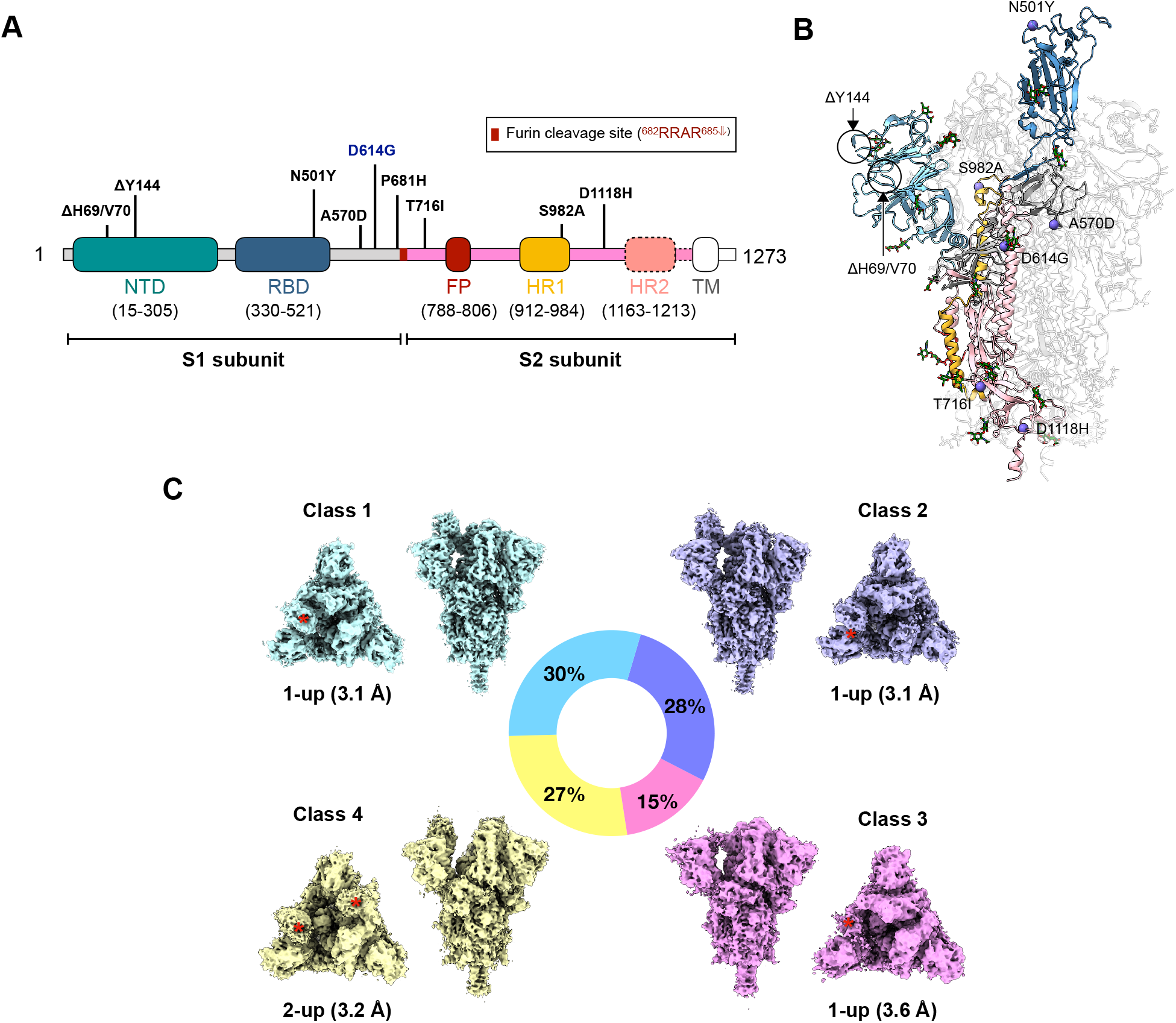
Cryo-EM structures of S-UK. (A) Domain structure of S-UK labeled with the nine B.1.1.7 mutations. The D614G mutation that is common in most emerging SARS-CoV-2 strains is highlighted in blue. (B) Four different classes of conformations of S-UK derived from cryo-EM 3DVA. The relative populations of the individual classes are indicated in the pie chart and the number of RBD in the upward conformation (RBD-up) of each class is indicated below the top view along with the nominal resolution in parenthesis. The positions of the RBD-up are indicated by red asterisks.

### Regulation of RBD orientation by a “pedal bin” mechanism unique to B.1.1.7

RBD-up conformation is the pre-requisite for ACE2 binding. Of all the reported cryo-EM structures of S-WT solved under physiological conditions (neutral pH), the majority exhibit an all RBD-down state with some in a single RBD-up state^16,21,22^. In contrast, the reported cryo-EM structures of S-D614G exhibit a higher propensity to populate RBD-up states with the majority being in one RBD-up state and some in two RBD-up state^17–20^. Our recent study indicated that none of the observed S-D614G particles were in an all RBD-down state, while 62 % of which had one RBD-up and 38 % had two RBD-up state^23^. Similar to S-D614G, S-UK did not populate all RBD-down state. Nevertheless, relative to S-D614G, S-UK exhibited less two RBD-up state (27 %) and predominantly populated the one RBD-up state (73 %) (Figure 1B). Gamblin and coworkers postulated that a salt bridge formed by D614 and K854 from two different protomers within the trimeric assembly is a linchpin to restrict inter-protomer motions to maintain RBD-down states. The D614G mutation abrogates the D614-K854 salt bridge thereby promoting RBD-up states^19,22^.

Close examination of the cryo-EM structure of S-UK indicated that the B.1.1.7-specific A570D mutation lies in close proximity to the D614G mutation site. The central helix in which K854 and K964 reside is flanked by D614G and A570D from the other protomer (Figure 2). The cryo-EM structure of S-UK in an RBD-up state exhibited a new salt bridge formed by K854_B_ and A570D_A_, accompanied by another salt bridge between K964_B_ and D571_A_ adjacent to A570D (Figure 2A, bottom panel). In contrast, the cryo-EM structure of S-UK in an RBD-down state exhibited an alternative salt bridge between K964C and A570DB (Figure 2A, top panel). These additional salt bridges likely compensate the loss of the salt bridge between D614 and K854 due to the D614G mutation. Comparison of the RBD-up and RBD-down conformations within the same S-UK structure revealed a 30° rotation of the RBD relative to the pivot point defined by the center of mass (COM) of the SD2 domain. In contrast, a minor rotation of the N-terminal domain (NTD) by 7° was observed (Figure 2B). The rigid body domain motion resulted in an upward displacement of 22 Å of the COM of RBD with a concomitant downward displacement of 3 Å of the COM of SD2 while the central helix remained static. Note that the downward displacement of SD2 was essentially governed by the downward loop movement promoted by the switch of salt bridge pairing that involves the A570D mutation. We propose that the switching of the A570D-mediated salt bridges may serve as the pedal of a pedal bin-like mechanism to modulate the RBD motion.

**Figure 2.**
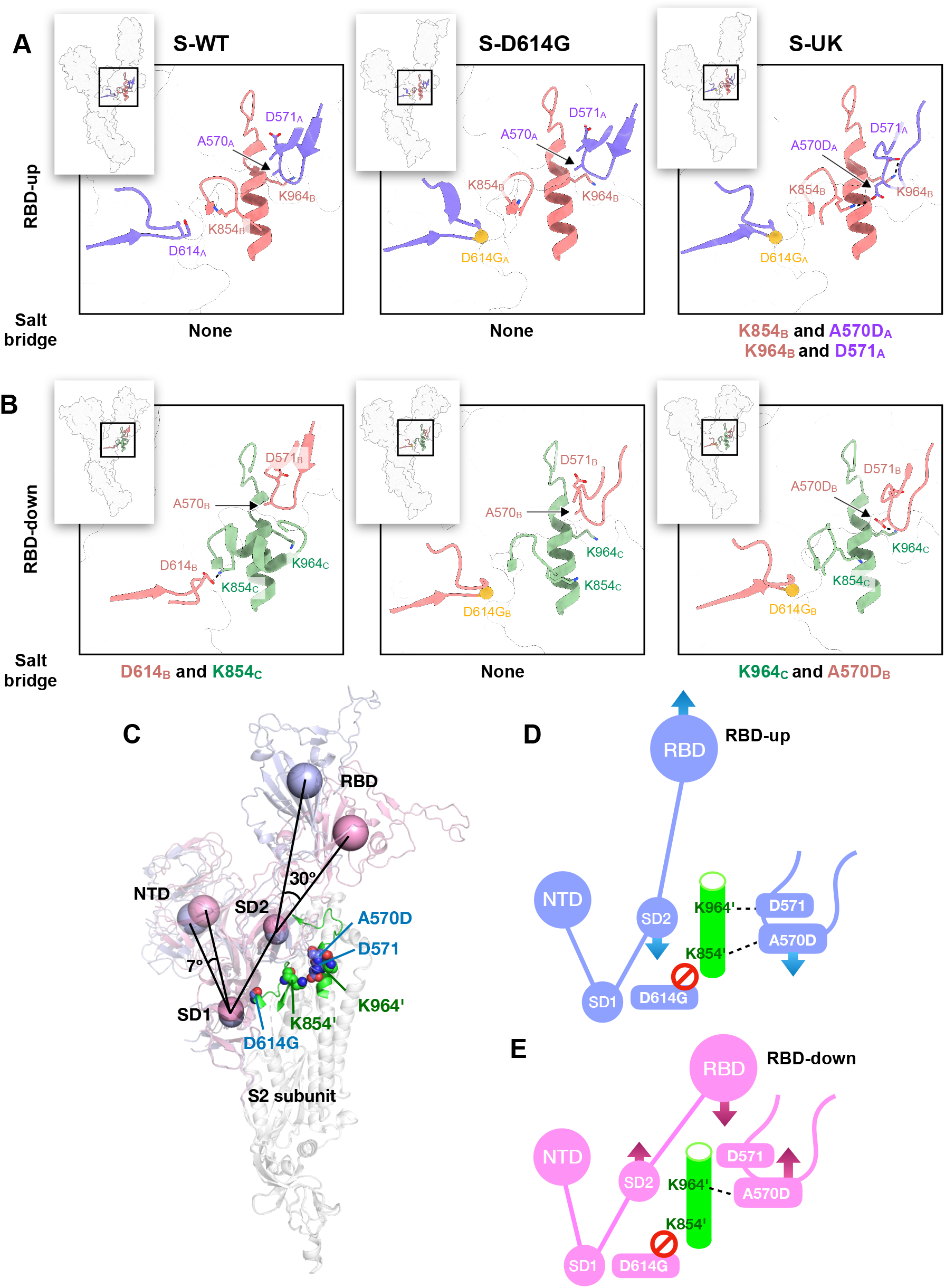
Molecular switch of regulating the RBD orientation. Structural basis of the interprotomer interaction networks around the clinical mutation sites for RBD-up (A) and RBD-down (B). Inset on the upper left corner of each panel illustrates the overall conformation of one protomer with the mutation site of interest indicated by a box, which is expanded in the corresponding panel. The RBD-up and RBD-down protomers are colored in light blue (chain A) and pink (chain B), respectively. For RBD-down, the corresponding protomer, chain B, is interacting with another RBD-down protomer, chain C, which is color in light green. (C) Comparison of the relative domain orientations of RBD-up and RBD-down in the context of the clinical mutations. The centers of mass of NTD (residues 1-288), RBD (residues 330-525), SD2 (residues 317-329 and 526-588), and SD1 (residues 290-316 and 589-695) are shown in spheres over the semitransparent cartoon representations of RBD-up (light blue) and RBD-down (pink). The two mutated residues, D614G and A570D, along with D571 on the same protomer, and K854 and K964 on the other protomer (indicated by an apostrophe) are shown in spheres. The alpha helix and the preceding loop in which K854’ and K964’ reside are shown in a cartoon representation and colored in green. Schematic representations of the domain motions of RBD-up (D) and RBD-down (E) modulated by the inter-protomer salt bridges. A switch in salt bridge pairing between K964’ and D571/A570D is proposed to regulate the pedal bin-like motions of RBD and Domain D.

### Molecular basis of ACE2 recognition by S-UK

To glean structural insights into the receptor binding of S-UK, we determined the cryo-EM structure of S-UK in complex with the ectodomain of ACE2 fused to a super-fold green fluorescence protein (hereafter ACE2). The 3DVA analysis of the complex demonstrated that all three RBDs of S-UK were bound to ACE2 in a 3:3 stoichiometry as opposed to the previously reported heterogeneous binding stoichiometry between the S protein of the Wuhan variant (hereafter S-WT) and ACE2 (Figure 3A-B and Supporting Information Fig. S3)^4,24,25^. Furthermore, the local orientation of the RBD in complex with ACE2 shows structurally dynamic, leading to the weak density and poor resolution (> 4.5 Å) of this region (Supporting Information Fig. S3). To this end, we therefore carried out the local refinement with a mask focusing on the RBD and ACE2 to result in a density map of 3.5 Å (Supporting Information Fig. S3). The improved resolution of the region favors us to confidently build the atomic model to uncover the binding interface between RBD and ACE2. Detailed inspection revealed a newly formed π-π stacking between the mutated Y501 of S-UK and Y41 of ACE2, which could potentially provide additional stability of the complex formation (Figure 3C). Furthermore, a contracted rigid-body domain movement of ACE2 towards the RBD of S-UK relative to that of S-WT by 2.5 Å was observed (defined by the COM displacement of ACE2).

**Figure 3.**
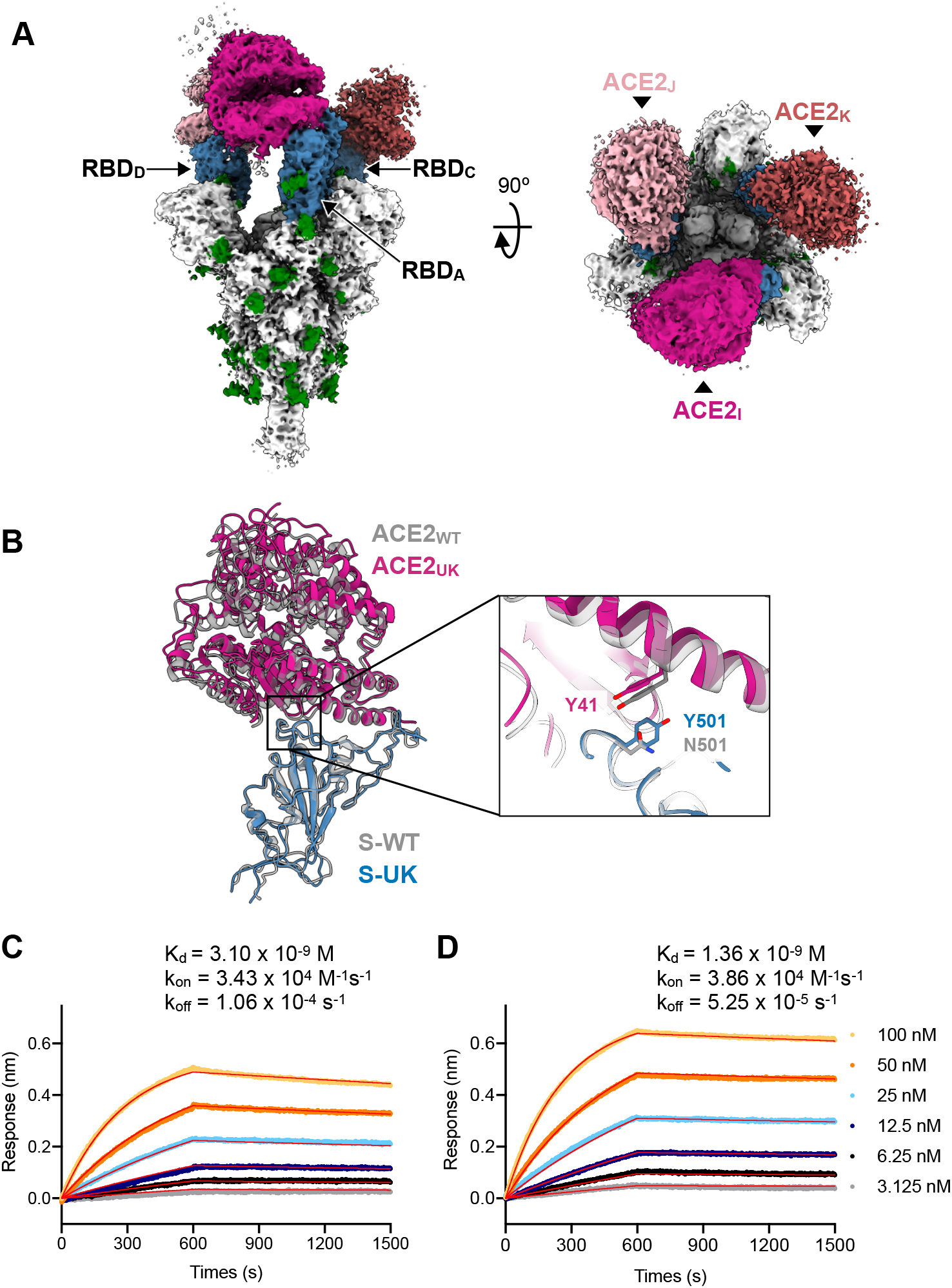
Structural basis and kinetics of ACE2 binding by S-UK. (A) Orthogonal views of the cryo-EM map of S-UK in complex with ACE2 in a 3:3 binding stoichiometry. (B) Expanded view of the atomic model of RBD in complex with ACE2. The RBD of S-UK is colored blue and S-UK-bound ACE2 is colored magenta. For comparison, the RBD of S-WT in complex with ACE2 is shown in grey. The N501Y mutation within the RBD introduced an additional pi-pi interaction with Y41 of ACE2 that could strength the intermolecular interaction. BLI sensorgrams of S-WT (C) and S-UK (D) binding to ACE2, which was immobilized onto the sensor tip. The concentrations of S-D614G and S-UK used in the independent BLI binding assays are indicated on the right. The kinetic parameters derived from global fitting of the sensorgrams are shown above with the fitted data shown in red lines that overlaid with the experimental curves.

To evaluate the impact of the B.1.1.7-specific mutations on the binding of ACE2, we immobilized ACE2 onto the sensor tips for biolayer interferometry (BLI) and compared the binding kinetics to S-UK and S-D614G. A two-fold decrease of the dissociation constant (K_d_) of S-UK (K_d_ = 1.3 nM) compared to that of the D614G variant (K_d_ = 3.1 nM) was observed (Figure 3D). The main difference was the reduced off-rate of S-UK (1.1×10^-4^ s^-1^) compared to that of S-D614G (5.3×10^-5^ s^-1^). Considering the slow dissociation kinetics of the binding event that could contribute to the uncertainty of the kinetic measurements, the effect of the N501Y mutation appeared to make marginal impact on the ACE2 binding kinetics.

### The N501Y mutation helps evade some but not all neutralizing antibodies

In spite of the marginal difference in ACE2 binding, the mutations in S-UK could help SARS-CoV-2 evade neutralizing antibodies to achieve higher infectivity^13^. To address this issue, we tested the abilities of three chimeric anti-RBD antibodies (RBD-chAbs) to compete with ACE2 binding to S-UK. We recently demonstrated that the three RBD-chAbs, namely RBD-chAb15, 25 and 45, targeted three distinct sites within the RBM, that each of these RBD-chAbs potently neutralize pseudovirus of the original Wuhan strain of SARS-CoV-2, and that the cocktail of RBD-chAb25 and 45 can prophylactically protect mice and hamsters from SARS-CoV-2 infection^26^. BLI analysis showed that pre-incubation of S-D614G with each of the three RBD-chAbs effectively prevented ACE2 binding. However, RBD-chAb25 failed to compete ACE2 binding to S-UK while RBD-chAb15 and 45 remained highly effective ACE2 binding inhibitors (Figure 4A-C). The loss of neutralizing activity of RBD-chAb25 against S-UK can be rationalized by the overlap of its structural epitope and the N501Y mutation evidenced by our recent cryo-EM analysis (Figure 3C and Supporting Information Fig. S4). Instead, epitope mapping of RBD-chAb15 and 45 reside on two distal ends of RBM thus potentially allowing simultaneous binding to the same RBD. To verify our hypothesis, we sequentially mixed RBD-chAb45 and 15 with S-D614G shortly before cryo-EM grid vitrification, and determined the cryo-EM structure of the quaternary complex (S-D614G:RBD-chAb15/45). The resulting cryo-EM map revealed stoichiometric binding of RBD-chAb15 and 45 to all three RBDs, and the N501 was bypassed by the two RBD-chAbs (Figure 4D-E). Despite the limited space above the three upward pointing RBDs to accommodate six antibodies, the poses of RBD binding by the two RBDs were essentially identical to that of individually bound cryo-EM structures, underscoring their potential to be used as a cocktail therapy for B.1.1.7.

**Figure 4.**
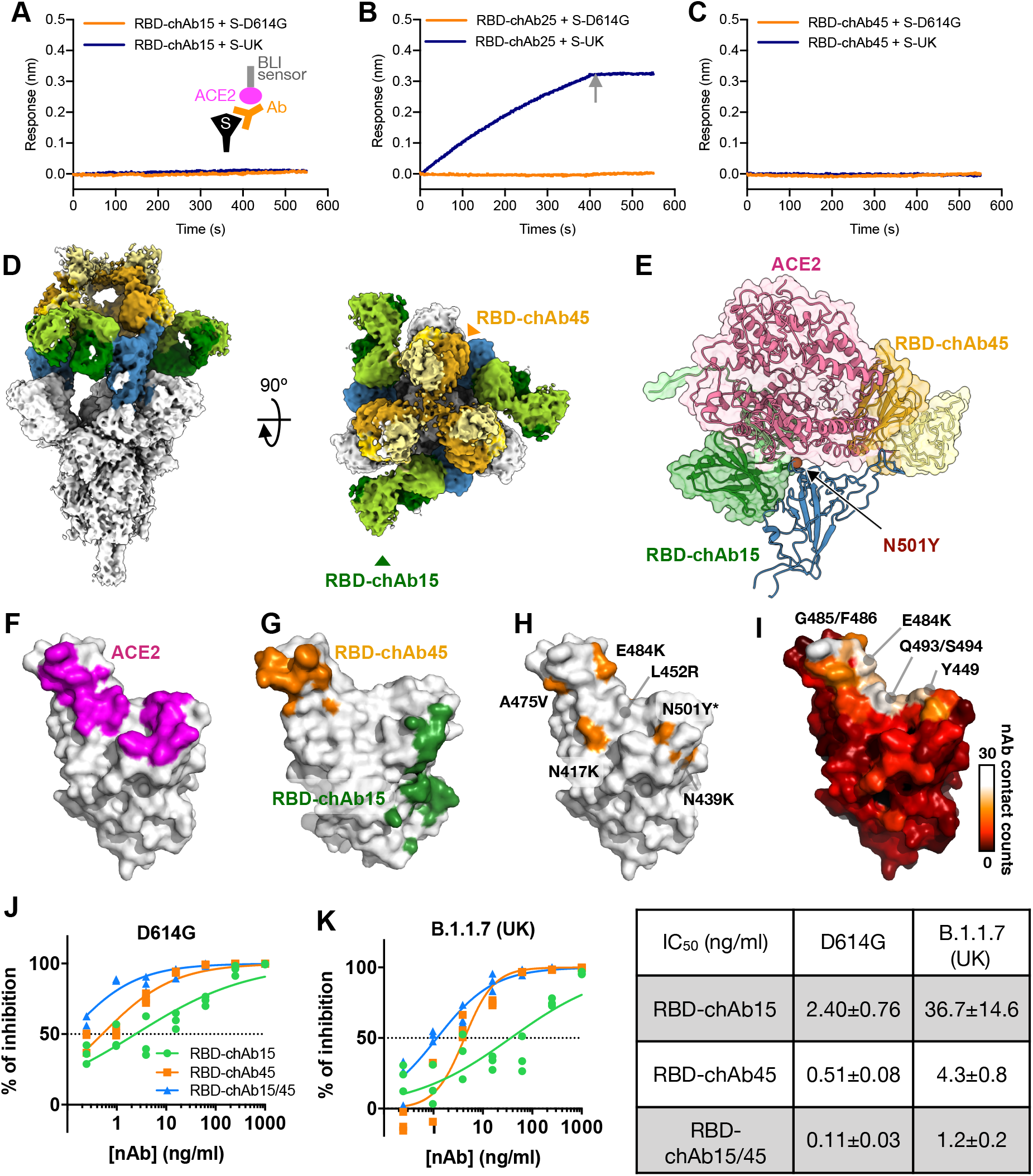
Neutralization of ACE2 binding by RBD-specific monoclonal antibodies. BLI sensorgrams of immobilized ACE2 binding to S-D614G and S-UK, both of which were preincubated with RBD-chAb15 (A), RBD-chAb25 (B), and RBD-chAb45 (C). The sensorgrams of S-D614G and S-UK are shown in orange and indigo, respectively. Only S-UK preincubated with RBD-chAb25 showed significant ACE2 binding, indicating the loss of neutralizing capacity of RBD-chAb25 as a result of the N501Y mutation. The grey arrow indicates the time point at which the dissociation of the complex was triggered. Schematic illustration of the experimental design is shown in (A). (D) Orthogonal views of the cryo-EM map of S-D614G in complex with RBD-chAb15 and 45 in a 3:3:3 binding stoichiometry. The three RBDs are shown in blue, the heavy chain and light chain of RBD-chAb15 in amber and yellow, respectively, the heavy chain and light chain of RBD-chAb45 in forest green and apple green, respectively. (E) Superposition of cryo-EM structure of RBD-bound ACE2 (PDB entry: 7KMS) ^4^ and that of the quaternary complex of antibody cocktail and RBD. The position of the N501Y mutation site is shown in a red sphere. (F) Structural mapping of ACE2 binding interface. The interface is defined as the atoms within RBD that are within 5 Å of ACE2, which is colored in magenta. (F) Structural mapping of antibody cocktail binding interface. RBD-chAb15 binding interface is colored in green. RBD-chAb45 binding interface is colored in orange. (H) Structural mapping of clinically reported mutations within RBD. The positions of the individual mutations are colored in orange and indicated with respective identities. N501Y, which is the only RBD mutation of S-UK is indicated by an asterisk. (I) Structural mapping of the contacting frequency of nAbs derived from convalescence sera. In total, 46 cryo-EM or crystal structures of nAb-bound RBD were analyzed by PISA to defined the intermolecular contacts. The frequency of nAb contacts is colored according to the heat map on the right. The identities of the most targeted residues are indicated. Pseudovirus analyses of D614G (J) and B.1.1.7 (K). The experiments were carried our in triplicate. The best-fit IC_50_ values and the corresponding standard errors of RBD-chAb15 and 45, and their combination derived from (J) and (K) are tabulated on the right.

To verify our hypothesis, we generated two pseudoviruses that individually expressed S-D614G and S-UK on the viral surface, and tested their pseudovirus neutralization activity of RBD-chAb15 and 45 in isolation and in combination (cocktail). The results showed that both nAbs were more effective against the D614G strain (Figure 4J) over the B.1.1.7 strain (Figure 4K), and that the cocktail of equal amounts of RBD-chAb15 and 45 was significantly more effective than the separately used nAbs.

## Discussion

In summary, we showed that the B.1.1.7-specific mutations minimally perturb the overall structure of the trimeric assembly of S-UK while a shift in the population of upward pointing RBD was observed. Compared to S-D614G, the two RBD-up population of S-UK was reduced without populating all RBD-down state. We identified a unique salt bridge switch involving the B.1.1.7-specific A570D mutation with two lysine residues, K854 and K964, in the central helix to compensate the loss of the salt bridge between D614 and K854 due to the D614G mutation. We postulate that the newly evolved A570D mutation serves as a molecular switch in a pedal bin mechanism to kinetically modulate the up/down motion of RBD. The RBD-up state is stabilized by the double salt bridge – A570D-K854 and D571-K964 – thereby increasing the kinetic barrier for the RBD-up to return to the RBD-down state relative to S-D614G. A recent report shows that introducing the A570D mutation to S-D614G leads to increased sensitivity of three RBD-up-specific antibodies, thus leading support to our proposed regulatory mechanism achieved by the A570D mutation^13^. Furthermore, our S-UK:ACE2 structure adopts a 3:3 binding stoichiometry unlike previously reported cryo-EM structures of ACE2-bound S-WT, the majority of which display a single RBD-up engaged with ACE2 binding^4,24,25^. This serves as additional evidence to support our conjecture that the A570D mutation helps stabilize the RBD-up state.

Increased RBD-up population helps host receptor binding at the expenses of increased probability of being targeted by nAbs. The N501Y mutation in S-UK introduces π-π staking between N501Y and Y41 of ACE2, according to our cryo-EM structure of S-UK. This translates into increased binding affinity between S-UK and ACE2 compared to that between S-D614G and ACE2 according to our BLI measurements. In the context of antibody neutralizing, S-UK harbors the N501Y mutation within RBM to disrupt binding by a subset of neutralizing antibodies, including RBD-chAb25 in our case. Nevertheless, RBD-chAb15 and 45 remained highly neutralizing in the context of competing ACE2 binding (Figure 4A-G). The quaternary cryo-EM structure of S-D614G:RBD-chAb15/45 showed that the two antibodies can simultaneously bind to the same RBD without contacting the N501Y mutation site. The pseudovirus assay further demonstrated that the combined use of RBD-chAb15 and 45 can synergistically neutralize pseudoviruses that correspond to D614G and B.1.1.7 (Figure 4J-K).

RBD-chAb45 targets the tip of the RBM where F486 resides. Statistical analysis of reported SARS-CoV-2 S protein bound to convalescence plasma-derived nAbs indicated that F486 is one of the most targeted RBD residues (Figure 4I). The adjacent E484K mutation that is present in the emerging South African 501.v2 strain and the Brazilian P1 strain could potentially reduce the efficacy of RBD-chAb45. Several point mutations are also reported to be partially overlapping with the high-frequency nAb epitopes, which could impede their neutralizing activities (Figure 4H). Nevertheless, the structural epitope of RBD-chAb15 is less frequently recognized by convalescence plasma-derived nAbs and there is hitherto no mutation in the more transmissible emerging SARS-CoV-2 variants that are located within the epitope of RBD-chAb15 (Figure 4G). Therefore, the ability of RBD-chAb15 and 45 to simultaneously bind to distinct regions of RBD is therefore an attractive feature for considering their use in prophylactic cocktail antibody therapy to prevent mutational viral escape.

## Methods

### Expression and purification of S-UK and S-D614G

The codon-optimized nucleotide sequence of S-UK harboring the double proline mutations (2P, ^986^KV^987^ → ^986^PP^987^) and the furin cleavage site mutation (fm, ^682^RRAR^685^ → ^682^GSAG^685^) for stabilizing the S protein in a prefusion state^27^ was a kind gift of Dr. Mi-Hua Tao (Institute of Biomedical Sciences, Academia Sinica). The DNA sequence corresponding the residues 1-1208 of S-UK was subcloned into the mammalian expression vector pcDNA3.4-TOPO (Invitrogen), which contains a foldon trimerization domain based on phage T4 fibritin followed by a c-myc epitope and a hexa-repeat histidine tag as described previously^28^. The same construct design was used for S-D614G as described elsewhere^23^.

The plasmids of S-D614G and S-UK were transiently transfected into HEK293 Freestyle cells for recombinant protein production as described previously^28^. The recombinant proteins were affinity purified by overnight binding to HisPur Cobalt Resin (Thermo Fisher Scientific, U. S. A.) in buffer A (50 mM Tris-HCl (pH 7.6), 300 mM NaCl, 5 mM imidazole and 0.02% NaN_3_ at 4°C), followed by washing with buffer B (50 mM Tris-HCl (pH 7.6), 300 mM NaCl, 10 mM imidazole), and elution by buffer C (50 mM Tris-HCl (pH 7.6), 150 mM NaCl, 150 mM imidazole). The eluted protein was further purified by size-exclusion chromatography (SEC) using a size-exclusion column (Superose 6 increase 10/300 GL; GE Healthcare, U. S. A.) in TN buffer (50 mM Tris-HCl (pH 7.6), 150 mM NaCl, 0.02 % NaN_3_). The protein concentrations were determined by using the UV absorbance at 280 nm using an UV-Vis spectrometer (Nano-photometer N60, IMPLEN, Germany).

### Expression and purification of sfGFP-ACE2

The sfGFP-ACE2 construct was obtained from Addgene (Plasmid #145171), which was deposited by Erik Proco (University of Illinois, USA)^29^. Recombinant sfGFP-ACE2 was transiently expressed in Expi293 cell and secreted into the culture medium. sfGFP-ACE2 was purified by an ion exchange column (HiTrap 5 ml Q FF anion exchange chromatography column; GE Healthcare, U. S. A.) eluted by a linear salt gradient from 10 to 1000 mM NaCl, followed by SEC using a size-exclusion column (Superdex 75 16/600 GL; GE Healthcare, U. S. A.) in TN buffer. The protein concentrations were determined by using the UV absorbance at 280 nm using an UV-Vis spectrometer (Nano-photometer N60, IMPLEN, Germany).

### Preparation of S-UK in complex with sfGFP-ACE2

Three microliters of S-UK in 2.3 mg/ml was mixed with 1.2 equivalent of sfGFP-ACE2 (S-UK:sfGFP-ACE2) and incubated at room temperature for 1 hr before being separated by SEC using a size-exclusion column (Superose 6 increase 10/300 GL; GE Healthcare, U. S. A.) in TN buffer. The elution peak corresponding to S-UK in complex with sfGFP-ACE2 was collected and concentrated to 1.5 mg/ml for cryo-EM grid preparation.

### Preparation of S-D614G in complex with RBD-chAb cocktail

Three microliters of S-D614G in 1.2 mg/ml was mixed with 1.2 equivalent of RBD-chAb45 and incubated at room temperature for 1 hr before being separated by SEC using a size-exclusion column (Superose 6 increase 10/300 GL; GE Healthcare, U. S. A.) in TN buffer. The elution peak corresponding to S-UK in complex with RBD-chAb45 was collected and concentrated to a total concentration of 1.5 mg/ml, followed by the addition of 1.2 equivalent RBD-chAb15 (S-UK:RBD-chAb15/45) for cryo-EM grid preparation. The preparation of RBD-chAb15 and 45 was described elsewhere^26^.

### Cryo-EM sample preparation

Three microliters of purified protein samples – S-UK, S-UK:sfGFP-ACE2, and S-UK:RBD-chAb15/45 – were applied onto 300-mesh Quantifoil R1.2/1.3 holyed carbon grids. The grids were glow-charged at 20 mA for 30 s. After 30 s incubation, the grids were blotted for 2.5 s under 4 °C with 100% humidity, and vitrified using a Vitrobot Mark IV (ThermoFisher Scientific, U. S. A.).

Data acquisition was performed on a 300 keV Titan Krios microscope equipped with a Gatan K3 direct detector (Gatan, U. S. A.) in the super-resolution mode using EPU software (ThermoFisher Scientific, U. S. A.). Movies were collected with a defocus range of −0.8 to −2.6 μm at a magnification of 81000 x, resulting in a pixel size of 0.55 Å. A total dose of 50 e^-^/Å^2^ was distributed over 50 frames with an exposure time of 1.8 second. The dataset was collected with an energy filter (slit width: 15-20 eV) and the dose rate was adjusted to 8 e^-^/pix/s.

### Image processing and 3D reconstruction

All 2x binned super-resolution raw movies were subject to Relion-3.0^30^ with dose-weighting and 5×5 patch-based alignment using GPU-based software MOTIONCOR2^31^. After motion correction, all corrected micrographs then were transferred to cryoSPARC v2.14^32^. Contrast transfer function (CTF) estimation was performed by patch-based CTF. The satisfied exposures, whose “CTF_fit_to_Res” parameters were between 2.5 and 4 Å, were selected and applied to particle picking. A small subset of micrographs was used for template-free blob picker. The picked particles were extracted with a box size of 384 pixels and followed by iterative rounds of 2D classification for filtering junk particles.

For S-UK, approximately 1.3 million particles were picked and classified by *ab-initio* reconstruction with C1 symmetry (class = 5), followed by heterogeneous refinement to generate five distinct classes (class = 5) and non-uniform refinement to generate an initial model. A mask was generated based on the initial 3D model for 3DVA by cryoSPARC v2 to generate five clusters of structural ensembles. Particle images corresponding to four of the five clusters were reextracted, un-binned, and non-uniformly refined with local CTF refinement to yield four cryo-EM maps with nominal resolutions ranging between 3.1 and 3.6 Å (Supporting Information Figure S1).

A similar imaging processing workflow was applied for S-UK:sfGFP-ACE2 and S-UK:RBD-chAb15/45. For S-UK:sfGFP-ACE2, five clusters of structural ensembles were also generated by 3DVA, but all particle images (n=463,191) were pooled and refined without symmetry constrain to yield a 3.3 Å cryo-EM map (Supporting Information Figure S3). The local resolution analysis was calculated using ResMap^33^. To improve the local resolution of ACE2 binding interface, a mask was created to include ACE2 and RBD for local refinement, which yielded a 3.5 Å cryo-EM of the subset of the S-UK:sfGFP-ACE2 structure. For S-UK:RBD-chAb15/45, approximately 2.3 million particles were picked for iterative 2D classification, which yield 78,790 particles for initial model building and templated particle picking, which generated 108,126 particles for 3D classification into three classes. The largest class (74%) was further refined by using a C3 symmetry to yield a 4.0 Å cryo-EM map (Supporting Information Figure S4). The local resolution analysis was calculated using ResMap^33^.

### Model building and refinement

An initial model of S-UK was generated based on PDB ID: 6XM3 by using Swiss-Model^34^. The coordinate was divided into individual domains and manually fitted into the cryoEM map in UCSF-Chimera^35^, UCSF-ChimeraX^36^ and Coot^37^. A similar approach was used for S-UK:sfGFP-ACE2, for which crystal structure of ACE2 in complex with RBD (PDB accession code 7KMS) was used as the initial model. For S-UK:RBD-chAb15/45, the previously determined structures of S-WT in complex with RBD-chAb15 and RBD-chAb45 were used as template for model building^26^. After iterative refinements, the coordinate was further processed by real-space refinement in Phenix^38^ to attain a convergent model. Based on the role of the N-glycosylation motif (N-X-S/T), N-linked glycans were built onto asparagine side-chains by using the extension module “Glyco” within Coot^37^. The final models were assessed by MolProbity^39^. Structural visualization and rendering of structural representations were achieved by using a combination of UCSF-Chimera, UCSF-ChimeraX and Pymol (Schrodinger Inc. U. S. A.).

### Biolayer interferometry (BLI)

A stock solution of sfGFP-ACE2 was diluted to at 10 μg/ml for immobilization onto High Precision Streptavidin (SAX) biosensors (Sartorius, Germany) in assay buffer (50 mM Tris-HCl, pH 7.6, 150 mM NaCl, 0.02% NaN_3_, 0.1% BSA) according to manufacture instructions. Stock solutions of S-D614G and S-UK were serially diluted to 100, 50, 25, 12.5, 6.25 and 3.125 nM in assay buffer for independent binding using an OctetRED 96 biolayer interferometer (ForteBio, USA). The association and dissociation were monitored over 600 s and 900 s, respectively. The six independent BLI sensorgrams with different S protein concentrations were baseline-corrected using double reference, i.e., biosensors without sfGFP-ACE2 were used to collect sensorgrams with another set of S protein samples to minimize non-specific binding and to extract the actual responses from the protein-protein interactions. The double reference subtracted data were globally fit to a 1:1 binding model using Data Analysis v10.0 software (ForteBio, USA). For RBD-chAb neutralization analysis, RBD-chAb15, 25 and 45 were individually mixed with an equal molar ratio of S-D614G or S-UK at a final concentration of 100 nM, and incubated at room temperature before BLI measurements using the same sfGFP-ACE2-immobilized biosensors with 400 s of association and 150 s of dissociation. The results were exported and replotted Prism 9 (GraphPad, USA).

### Structure-based statistics of RBD epitope binding frequency

The identification of structural epitopes of convalescent sera-derived RBD-specific nAbs at a residue-specific level as achieved by using the ‘Protein interfaces, surfaces and assemblies’ service PISA at the European Bioinformatics Institute (http://www.ebi.ac.uk/pdbe/prot_int/pistart.html)^40^. The list of cryo-EM and crystal structures of nAb-bound spike and RBD was manually curated by searching the entries deposited in the Protein Data Bank and their corresponding publications as tabulated in Tables S3-S4. The nAb contacting residues within RBD were defined by the function “Interfaces” of PISA and summarized in Table S5. In cases where one RBD residue is in contact with both the heavy chain and a light chain of a given nAb, the epitope count was considered to be twice. The cumulated epitope counts were mapped onto the crystal structure of RBD in complex with ACE2 (PDB accession code 6M0J) and rendered by using Pymol (Schrodinger Inc. U. S. A.).

### Pseudovirus neutralization assay

The pseudovirus neutralization assays were performed using 293T cells overexpressing ACE2 and pseudoviruses expressing full-length S protein provided by RNAi Core of Academic Sinica. Four-fold serially dilution of chAbs were premixed with 1000 TU/well D614G, UK or SA strain of SARS-CoV-2 pseudovirus. The mixture was incubated for 1 h at 37°C and then added to preseeded 293T-ACE2 cells at 100 μl/well for 24 h at 37°C. The medium was removed and refilled with 100 μl/well DMEM for additional 48-h incubation. Next, 100 μl ONE-Glo™ luciferase reagent (Promega) was added to each well for 3-min incubation at 37°C. The luciferase activities were measured using a microplate spectrophotometer (Molecular Devices). The inhibition rate was calculated by comparing the luminescence value to the negative and positive control wells. IC50 was determined by a four-parameter logistic regression using GraphPad Prism (GraphPad Software Inc.).

## Associated content

Experimental materials and methods, including molecular cloning, protein expression and purification, data collection and analyses of cryo-EM, BLI and pseudovirus data. The atomic coordinates of the S-UK are deposited in the Protein Data Bank (PDB) under the accession codes 7EDF, 7EDG, 7EDH, and 7EDI. Those of S-UK:ACE2 and S-D614G:RBD-chAb15/45 are deposited in the PDB under the accession codes 7EDJ and 7EH5, respectively. The cryo-EM maps are deposited in the Electron Microscopy Data Bank (EMDB) under successive codes from EMD-31069 to EMD-31074.

## Supporting Information

The supporting information includes:

Figures S1-S4, Table S1-S4 and references.

## Author contributions

S.-T.D.H. and H.C.W. conceived the experiment. T.J.Y. and P.Y.Y. prepared the spike proteins and ACE2. W.Y.C and H.T.L. generated and prepared RBD-chAbs. T.J.Y. and Y.C.C. collected the cryo-EM data. T.J.Y. processed the cryo-EM data and modeled the structures. T.J.Y. and M.R.H. collected and analyzed BLI data. K.H.L and H.C.T. collected and analyzed pseudovirus data. P.Y.Y. curated the RBD epitope usage data. T.J.Y. and S.-T.D.H. wrote the manuscript with inputs from all co-authors. S.-T.D.H. and H.C.W. obtained funding.

## Acknowledgement

We thank the Academia Sinica Biophysics Core Facility (Grant AS-CFII108-111), and Academia Sinica Cryo-EM Center (Grant AS-CFII-108-110) for data collection, all of which are funded by the Academia Sinica Core Facility and Innovative Instrument Project. We also thank the mammalian cell culture facility of Institute of Biological Chemistry, Academia Sinica, for supporting the protein production, and the Academia Sinica Grid Computing for cryo-EM data processing.

