## Supporting information for "Impacts on the structure-function relationship of SARS-CoV-2 spike by B.1.1.7 mutations"

**The supporting information includes:**

Fig. S1-S4, Table S1-S4 and references.

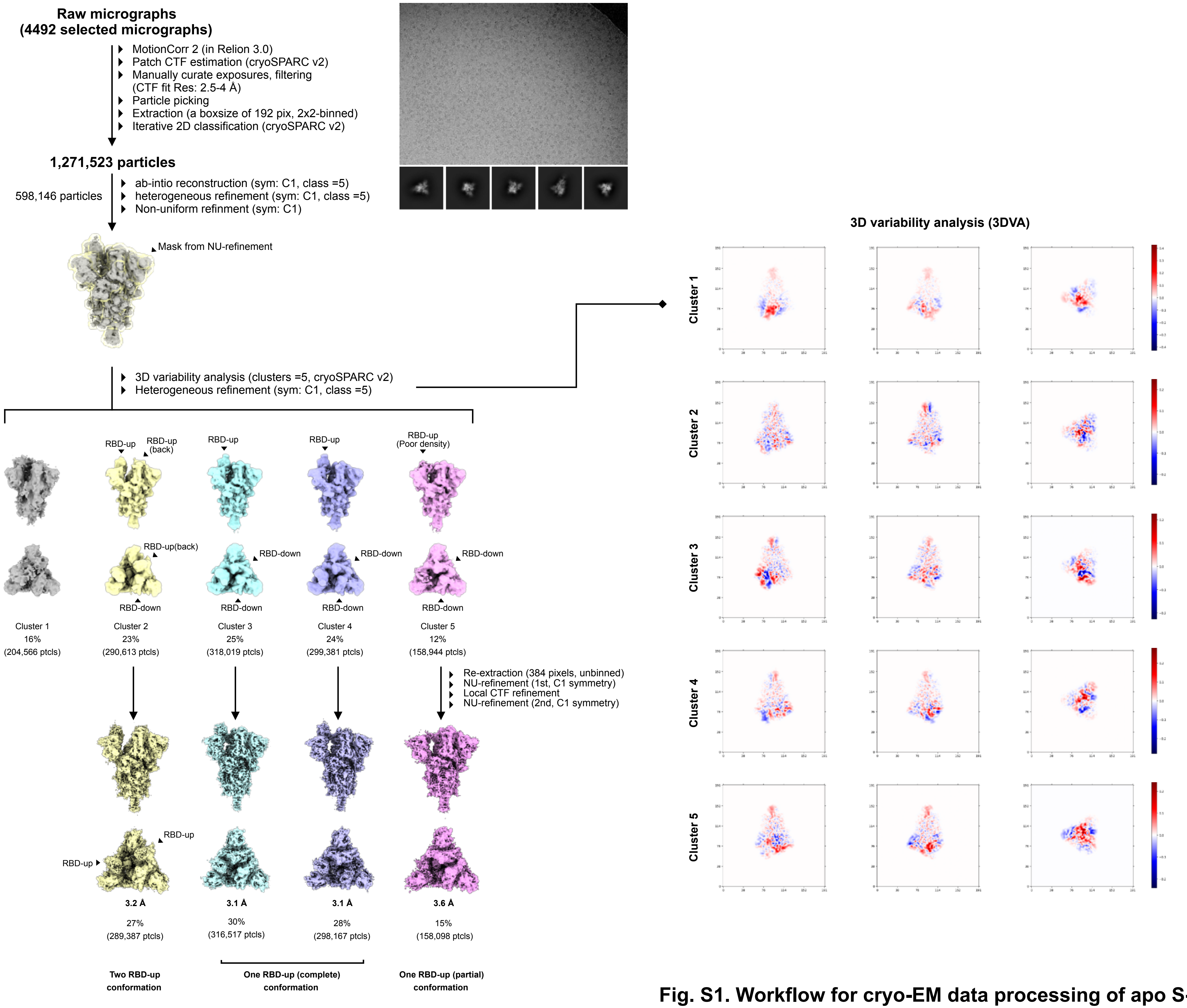


**Fig. S1. Workflow for cryo-EM data processing of apo S-UK**

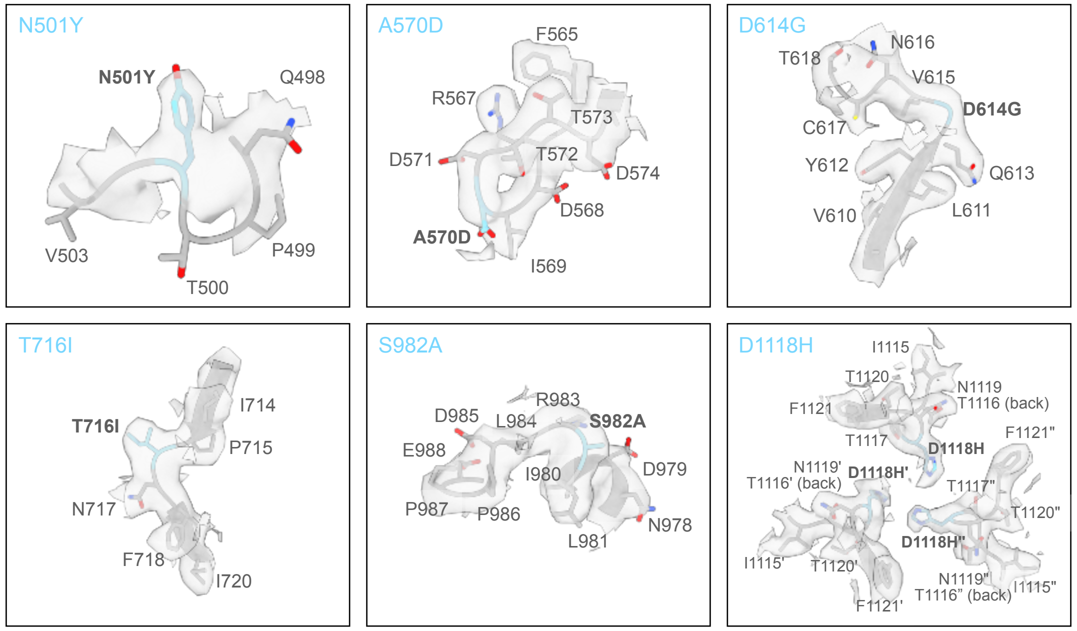


**Fig. S2. Expanded views of cryo-EM maps and atomic models of apo S-UK at the UK variant-specific mutations sites**

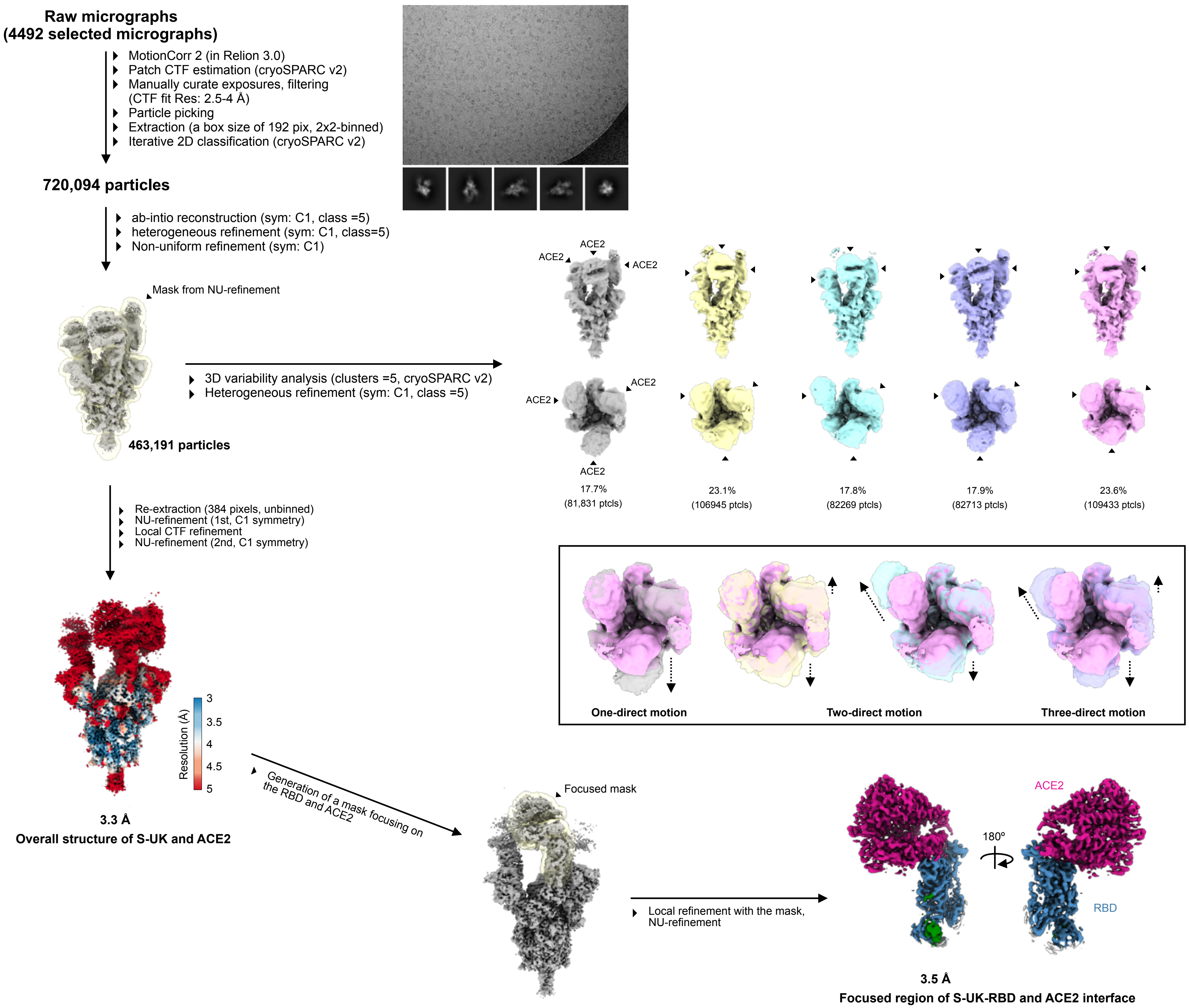


**Fig. S3. Workflow for cryo-EM data processing of S-UK:ACE2**

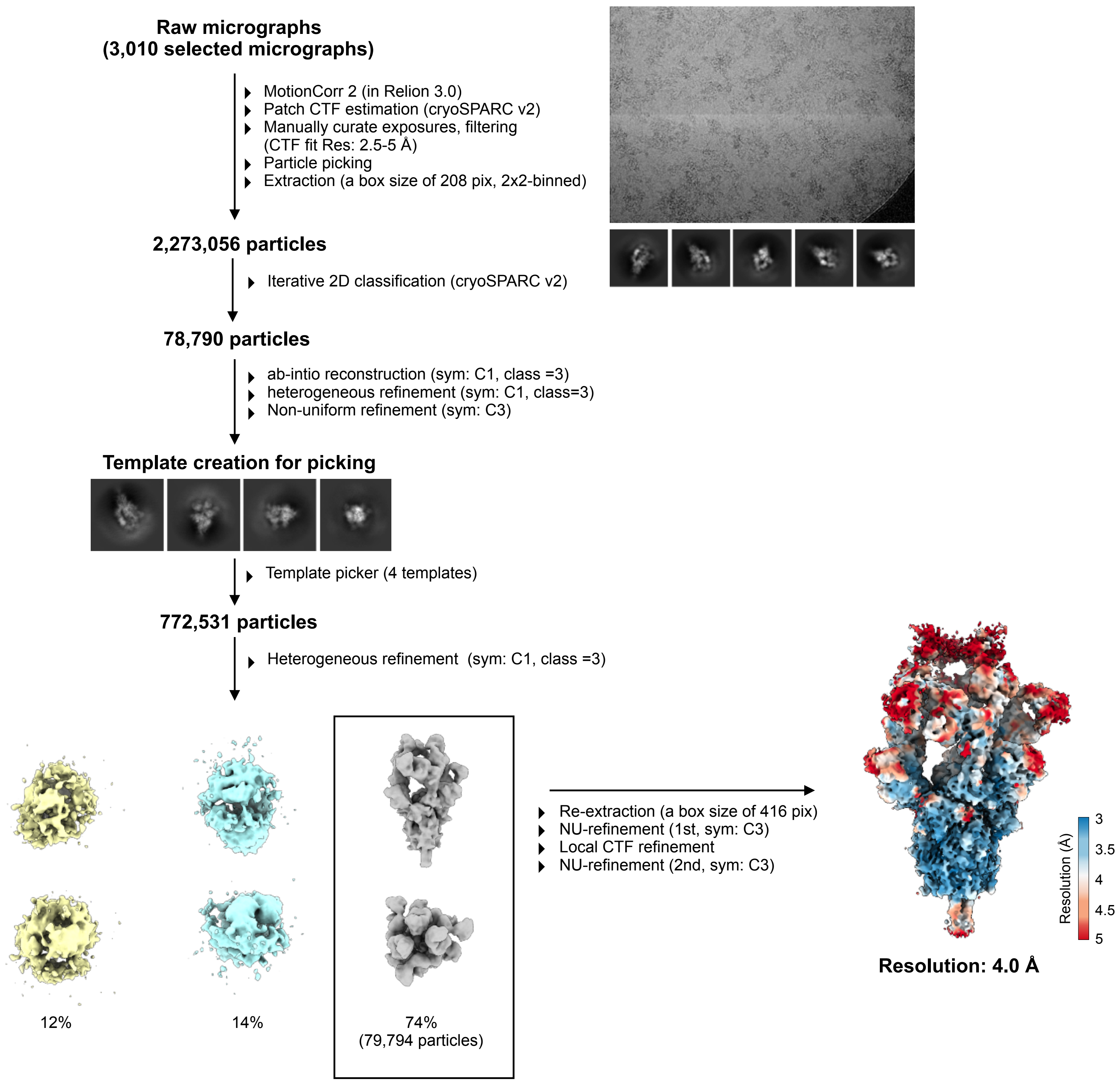


**Fig. S4. Workflow for cryo-EM data processing of S-D614G:RBD-chAb15/45**

**Table S1. Parameters of cryo-EM data collection, processing and model validation (apo S-UK).**

| | One RBD up conformation 1 | One RBD up conformation 2 | One RBD up conformation 3 | Two RBD up conformation 1 |
|---|---|---|---|---|
| **Data collection and processing** | | | | |
| Microscope | Titan Krios (Gatan K3 Summit camera) | | | |
| Voltage (keV) | 300 | | | |
| Mode | Counting | | | |
| Magnification | 81,000x | | | |
| Total dose ($e^{-}$/Å$^{2}$) | 50 | | | |
| Defocus range (μm) | 0.8-2.6 | | | |
| Pixel size (Å) | 1.1 (2x binned) | | | |
| Final used particles | 316,517 | 298,167 | 289,387 | 158,098 |
| Symmetry | C1 | C1 | C1 | C1 |
| Map Resolution (Å) | 3.2 | 3.2 | 3.6 | 3.3 |
| **Model composition** | | | | |
| Non-hydrogen atoms | 25,093 | 24,718 | 23,279 | 24,989 |
| Protein residues | 3,065 | 3,023 | 2846 | 3,048 |
| Ligands | 82 | 79 | 77 | 78 |
| MolProbity score | 1.75 | 1.79 | 1.81 | 1.86 |
| *Ramachandran (%)* | | | | |
| Favored | 93.87 | 92.99 | 93.34 | 93.40 |
| Allowed | 6.00 | 6.77 | 6.37 | 6.10 |
| Outliners | 0.13 | 0.24 | 0.29 | 0.50 |
| Rotamer outliners (%) | 0.19 | 0.19 | 0.16 | 0.19 |
| Clashscore | 6.23 | 6.20 | 6.92 | 7.94 |
| *r.m.s. deviations* | | | | |
| Bond length (Å) | 0.003 | 0.003 | 0.003 | 0.003 |
| Bond angles (º) | 0.643 | 0.607 | 0.602 | 0.640 |
| **Data Deposition** | | | | |
| PDB code | 7EDF | 7EDG | 7EDH | 7EDI |
| EMDB code | 31069 | 31070 | 31071 | 31072 |

**Table S2. Parameters of cryo-EM data collection, processing and model validation (S-UK/ACE2 complex and nAb cocktails).**

| | S-UK:ACE2 | S-D614G:RBD-chAb15/45 |
|---|---|---|
| **Data collection and processing** | | |
| Microscope | Titan Krios (Gatan K3 Summit camera) | |
| Voltage (keV) | 300 | |
| Mode | Counting | |
| Magnification | 81,000x | |
| Total dose ($e^-/Å^2$) | 50 | |
| Defocus range (μm) | 0.8-2.6 | |
| Pixel size (Å) | 1.1 (2x binned) | |
| Final used particles | 463,191 | 79,794 |
| Symmetry | C1 | C3 |
| Map Resolution (Å) | 3.3 | 4.0 |
| **Model composition** | | |
| Non-hydrogen atoms | 39,571 | 34,992 |
| Protein residues | 4,836 | 4,362 |
| Ligands | 84 | 75 |
| MolProbity score | 1.95 | 1.86 |
| *Ramachandran (%)* | | |
| Favored | 91.84 | 94.36 |
| Allowed | 7.99 | 5.50 |
| Outliners | 0.17 | 0.14 |
| Rotamer outliners (%) | 0.00 | 0.00 |
| Clashscore | 8.42 | 8.91 |
| *r.m.s. deviations* | | |
| Bond length (Å) | 0.005 | 0.002 |
| Bond angles (º) | 0.659 | 0.515 |
| **Data Deposition** | | |
| PDB code | 7EDJ | 7EH5 |
| EMDB code | 31073 | 31074 |

**Table S3. Cryo-EM structures used for RBD epitope counting.**

| Antibody | PDB code | EMDB | Reference |
|---|---|---|---|
| S2H13 | 7JV6 | 22494 | [1] |
| S2H13 | 7JV4 | 22492 | [1] |
| S2A4 | 7JVC | 22506 | [1] |
| S2M11 | 7K43 | 22659 | [2] |
| S2E12 | 7K4N | 22668 | [2] |
| C104 | 7K8U | 22731 | [3] |
| C119 | 7K8W | 22733 | [3] |
| C002 | 7K8T | 22730 | [3] |
| C104 | 7K8V | 22732 | [3] |
| C135 | 7K8Z | 22736 | [3] |
| C121 | 7K8X | 22734 | [3] |
| C121 | 7K8Y | 22735 | [3] |
| C144 | 7K90 | 22737 | [3] |
| C002 | 7K8S | 22729 | [3] |
| BD-308-2 | 7CHH | 30374 | [4] |
| 2-4 | 6XEY | 22156 | [5] |
| C105 | 6XCN | 22128 | [5] |
| C105 | 6XCM | 22127 | [5] |
| EY6A | 6ZDH | 11174 | [6] |
| Ab23 | 7BYR | 30247 | [7] |

**Table S4. Crystal structures used for RBD epitope counting.**

| Antibody | PDB code | Reference |
|---|---|---|
| CV07-250 | 6XKQ | [8] |
| CV07-250 | 6XKP | [8] |
| S2A4 | 7JVA | [1] |
| S2H13 | 7JV2 | [1] |
| S2E12 | 7K45 | [2] |
| BD604 | 7CH4 | [4] |
| BD629 | 7CH5 | [4] |
| BD604 | 7CHF | [4] |
| BD368-2 | 7CHF | [4] |
| BD-236 | 7CHE | [4] |
| BD-368-2 | 7CHE | [4] |
| BD-236 | 7CHB | [4] |
| BD-629 Fab/BD-368 Fab | 7CHC | [4] |
| BD-368-2 Fab | 7CHC | [4] |
| COVA2-39 | 7JMP | [9] |
| COVA2-04 | 7JMO | [9] |
| CC12.1 | 6XC2 | [10] |
| CC12.3 | 6XC4 | [10] |
| CR3022 | 6XC3 | [10] |
| CC12.1 | 6XC3 | [10] |
| CR3022 | 6XC7 | [10] |
| CC12.3 | 6XC7 | [10] |
| EY6A | 6ZFO | [6] |
| CV30 Fab | 6XE1 | [11] |
| COVA1-16 | 7JMW | [12] |
| B38 | 7BZ5 | [13] |
